# Evidence in favor of abrupt over gradual learning in the differential reinforcement of response duration (DRRD) task

**DOI:** 10.64898/2025.12.26.696617

**Authors:** Mateus Gonzalez de Freitas Pinto, Chang Chiann, Alexei Magalhães Veneziani, Marcelo Bussotti Reyes

## Abstract

Learning can occur in markedly different ways: in some cases, it unfolds as a gradual process, with behavior improving slowly toward an asymptotic level of performance; in others, it appears as an abrupt process that sharply separates behavior before and after a change point. Understanding the behavioral and neural processes underlying these distinct acquisition patterns may be critical for elucidating the basic principles of learning. We investigated this question experimentally using naïve rats performing a differential reinforcement of response duration (DRRD) task, in which animals were required to remain inside a nosepoke for a minimum duration of 1.5 seconds to get a sugar pellet as a reward. All rats learned to wait longer in the nosepoke when comparing behavior at the beginning and at the end of the experiment. We tested several continuous models against a single change point (CP) model, in which behavior changes at a specific moment and remains stable thereafter. Instead of the traditional approach based on trial-segmented behavior, we used the real time elapsed since the beginning of the experiment as a continuous, uncontrolled variable. We fitted all models to data from individual rats and compared model fit quality across alternatives. Our results provide strong evidence in favor of an abrupt change, as captured by the CP model, over all other models. Moreover, the residuals of the CP model exhibited a Gaussian distribution, suggesting that no additional systematic dynamics remained unexplained and that the behavioral dynamics were fully captured by a single change point.

## 1. Introduction

Is learning a gradual, continuous process that improves with each experience, or is it a sudden transition that occurs in a single decisive moment? This fundamental question has long been a subject of debate among researchers across the domains of psychology and neuroscience [14]. Some forms of acquisition clearly build on a single experience, as in declarative memory experiments with humans or fear conditioning in other animals, for example. These forms of acquisition are often referred to as “single-trial”, “one-shot” learning [1], or “eureka” [2], and predict abrupt shifts in behavior following a critical event. Other types of acquisition seem to evolve gradually over time, such as in appetitive classical conditioning tasks or procedural skill acquisition, as classically described by Rescorla and Wagner[19]. Disentangling these two possibilities is particularly challenging in tasks where behavioral variability and intrinsic noise mask the underlying learning dynamics. One such task is the **Differential** Reinforcement of Response Duration (DRRD) paradigm.

In the DRRD task, animals are required to produce responses (e.g., pressing a lever or nose-poking) that must exceed a fixed temporal threshold—referred to as the criterion—to obtain reinforcement. Responses shorter than this threshold typically result in a timeout, during which further responses are not reinforced, or rats are impeded from responding. This task has been used to study temporal learning, impulsivity, and behavioral inhibition. However, DRRD studies have traditionally focused on the analysis of trial-by-trial response durations, virtually ignoring the temporal pacing in which these responses occur. For instance, inter-trial intervals (ITIs)—the pauses between successive responses—are typically ignored, despite containing valuable information about motivation, timing strategy, and learning dynamics. Furthermore, a substantial body of evidence shows that ITI critically influences learning, at least in some contexts [24, 4, 3, 12].

In this study, we follow an alternative path to the traditional trial-based analysis and approach the subject by considering the animal’s behavior as a continuous process unfolding over the entire session time. By abandoning the assumption that learning depends on discrete trials, we aim to recover temporal patterns that might be obscured when experimenters segment behavior into predefined (perhaps artificial) units. This perspective allows for an alternative way to investigate whether learning in the DRRD task occurs smoothly over time or whether it reflects a more abrupt, step-like process [21, 11, 20, 13].

To address this question, we use recent advances in time series modeling for inhomogeneous data—data collected at irregular time intervals. Techniques originally developed in domains such as high-frequency finance [7, 6] and astronomy [9, 10, 8, 17] have proven useful for analyzing behavioral datasets with uneven sampling, such as those observed in free-operant conditioning paradigms. In particular, we adopt statistical tools that model the behavior over continuous session time, accounting for the irregular sampling inherent in DRRD tasks.

Inspired by methods such as the Lomb-Scargle periodogram [16, 22] and by approaches rooted in the Wiener-Khintchine theorem [25, 15], we examine whether the temporal evolution of response behavior exhibits characteristics consistent with smooth adaptation or whether sharp transitions better explain the data. By comparing model fits that assume gradual change with those that allow for abrupt shifts, we aim to provide empirical insight into the learning dynamics of interval timing.

## 2. Materials and methods

The data underlying the results reported in this article are publicly available through the Open Science Framework at osf.io/uh4s6.

### Subjects

Eight adult male Wistar rats (12–16 weeks old, 350–400 g mass) were used. All animals were naive to the experimental procedures and maintained under a 12:12 h light/dark cycle (lights on at 7:00 a.m.). They were housed individually with *ad libitum* access to water and restricted access to food to maintain 85% of their free-feeding body weight. All procedures were approved by the Institutional Animal Care and Use Committee of UFABC (CEUA-UFABC, permits 001/2014 and 020/2015). Additional procedural details are available in Tunes et al. [23].

### Apparatus

Behavioral training took place in custom-built acrylic operant chambers controlled by Arduino microcontrollers. Each chamber featured an infrared-based nose poke detector and an automated sucrose delivery system. A stepper motor controlled a gate that provided access to a metallic nozzle delivering a 50% sucrose solution. An electrostatic sensor monitored licks and triggered the gate to close after three licks.

### Behavioral Task

The behavioral paradigm was a self-initiated Differential Reinforcement of Response Duration (DRRD) task. Prior to timing training, rats were shaped using a fixed ratio 1 (FR1) schedule, in which each nose poke or lever press immediately resulted in reward access. Rats advanced to the DRRD phase after completing at least 100 rewarded responses in a single 60-minute session. All animals completed two DRRD sessions longer than 2.5 hours.

In the DRRD task, each trial was initiated by the animal entering the nose poke port. To receive a reward, rats were required to maintain their nose poke for a minimum of 1.5 seconds. Shorter responses resulted in no reward, and a new trial could begin immediately. For reinforced trials, rats gained access to three licks of a 50% sucrose solution. For each trial, both response duration and intertrial interval (ITI)—defined as the time between exiting and re-entering the nose poke—were recorded. Although there was no upper limit on response duration, more than 94% of all responses were shorter than 3.5 seconds.

### Data

Raw data consisted of time-stamped event codes indicating when the animal entered or exited the nose poke, as well as when the sucrose-delivery gate began opening and closing. This information was segmented into individual trials, allowing us to determine the precise start and end times of each response. Unlike traditional trial-based averaging methods, response durations were analyzed as a function of elapsed time since the beginning of the session.

Figure 1 depicts eight time-dependent signals representing in the Y-axis the durations of lever presses by different rats, numbered out from 3 to 10. The X-axis indicates the timing of each lever press, providing a temporal context for the observed behaviors.

**Figure 1:**
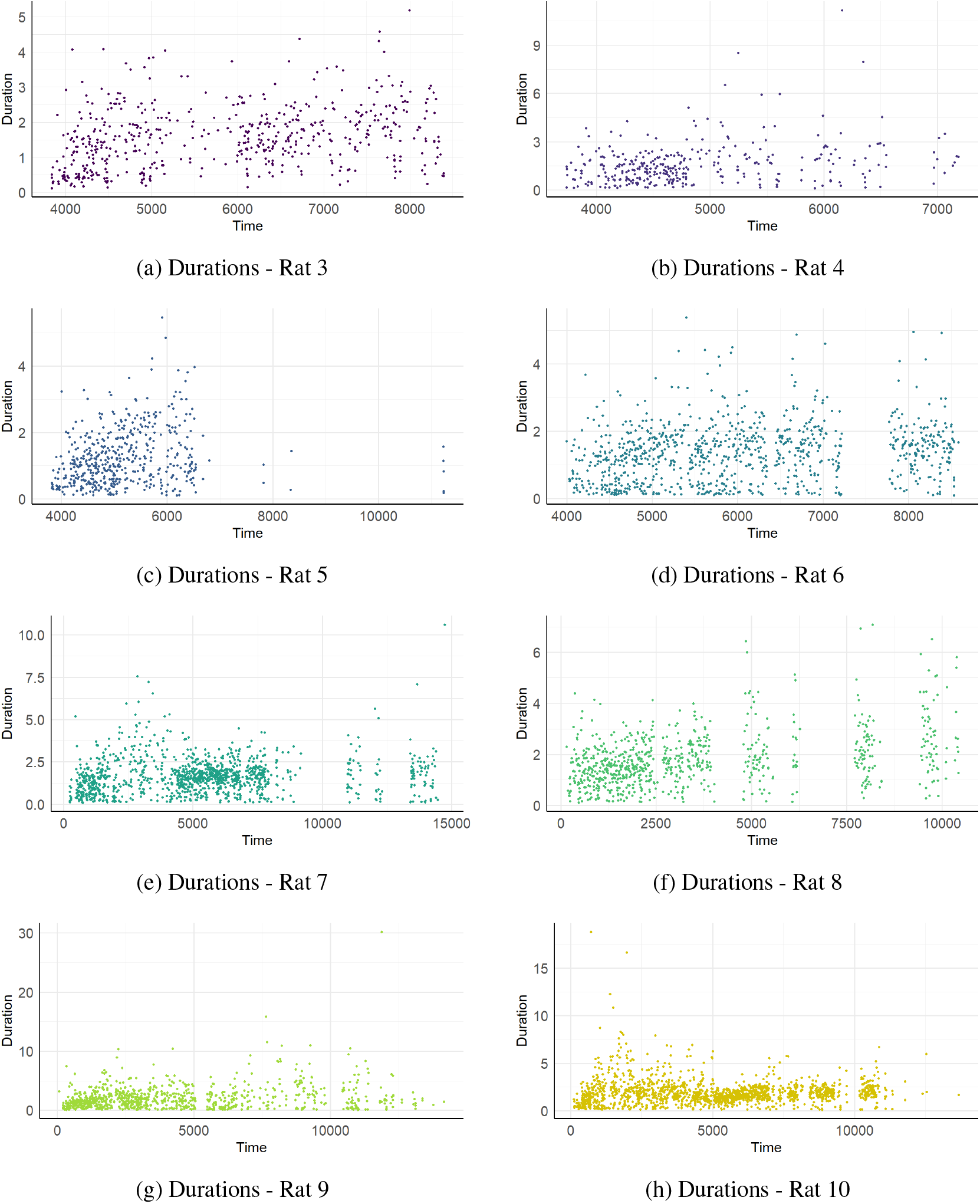
Durations for rats

As observed in Figure 1, the intervals between consecutive button presses vary, resulting in events that are not uniformly spaced over time. This lack of temporal regularity renders traditional time series analysis methods unsuitable, as they typically require equally spaced observations. Consequently, alternative techniques specifically designed for unevenly spaced time data must be employed.

It is also possible to see that rats show bursts of activity interspersed with quieter periods, suggesting alternating phases of engagement and rest. The duration spread (Y-axis) in Figure 1 differs markedly between rats: while some exhibit high-amplitude outliers, implying occasional unusual long presses, others show more compact distributions around shorter durations. This het-erogeneity may reflect the idiosyncrasies of each rat or differences in motor control, motivation, or fatigue.

Despite a similar task, the individual panels in Figure 1 suggest that the response dynamics is not straightforwardly uniform between subjects, with rats sharing some patterns, although with some variegated behavior. In particular, what is visually shared among some rats (3, 5, 7 and 9) is a high density of shorter durations at the beginning of the experiment, followed by periods where duration appears slightly higher. For other rats (6 and 8), it is not clear whether a single change point could better explain the data or whether a linear trend is a better fit. Nevertheless, this shared behavior of apparent shift in means is fundamental for our analysis of learning effects, which will be explored in further sections.

#### 2.1. Statistical methodology

We investigated a range of statistical models to characterize the temporal dynamics of the data and to enhance understanding of the underlying learning processes. As a starting point, we consider linear regression models and their functional variants, including logarithmic linear, exponential, and square root transformations. These formulations align with the classical theoretical conceptions of learning as a gradual and continuous process [19]. To capture potential periodicity in the data, we extend the linear model to include harmonic terms, considering *K* = 4 harmonics in the regression. This allows for the modeling of periodic components, which may reflect cyclic or repetitive structures in the learning trajectory.

In addition to these parametric models, we assess the performance of the irregular autoregressive model of order 1 (iAR(1)), previously applied in astrostatistics to account for irregular and inhomogeneous data structures [9, 10, 8, 17, 18]. We examine several variants of the iAR(1) model, including those with normally distributed residuals, gamma-distributed residuals, and normally distributed residuals combined with a linear trend component.

Moreover, we explore the hypothesis suggested in the literature that learning may occur abruptly or discontinuously, similar to a “snap” or phase transition [11, 20]. To model this, we include a change-point analysis in the mean structure of the data. Specifically, we tested for a single change-point *T* ^*^, with distinct mean levels *µ*_1_ for *t* ≤ *T* ^*^ and *µ*_2_ for *t* > *T* ^*^. Finally, we integrate this change-point structure into the iAR(1) framework by modeling it as a change in the underlying trend.

More formally, let *y*_*t*_ denote the observed result (trial duration) in continuous time *t* > 0. The following linear models are considered:

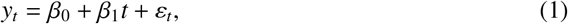

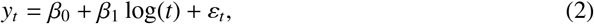

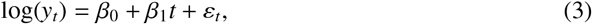

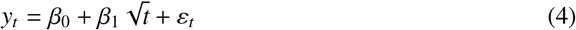

We hereinafter refer to models (1)-(4), respectively, as linear model, log-linear model, exponential-linear model, and square-root linear model. In all cases, the error term is assumed to follow a Gaussian distribution with unknown variance, which means *ε*_*t*_ ∼ *N*(0, *σ*^2^).

To capture possible periodicity in the learning process, we model the signal using trigonometric components:

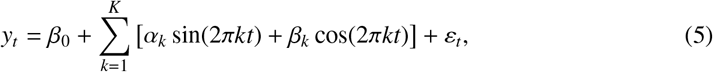

where in practice we set *K* = 4 harmonics and ε_*t*_ ∼ *N*(0, *σ*^2^).

Since observations of durations occur at irregular times *t*_1_ < *t*_2_ < · · · < *t*_*n*_, the usual ARMA models of [5] are not suitable, as they assume regularly spaced observations. Instead, we employ the iAR(1) model, given by:

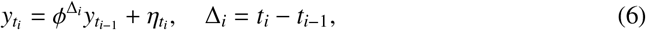

where 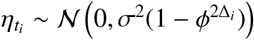, and *ϕ* ∈ (0, 1) captures the dependence across irregular intervals.

We can model the observed series as the sum of a deterministic trend 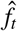 (obtained, for instance, from the exponential or log-linear models above) and a residual iAR(1) process:

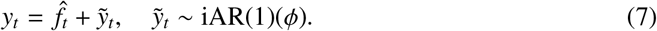

To account for asymmetric and positive-definite residuals, we also consider the iAR(1)-Gamma model in place of the usual Gaussian noise term.

Finally, we test the hypothesis that the mean of the process shifts at a single change-point *T* ^*^:

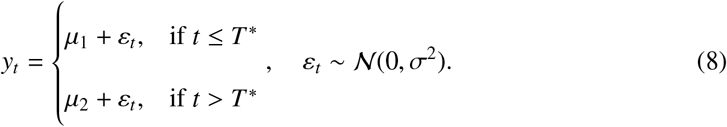

We extend the iAR(1) model to include autocorrelation with a structural break in the mean as:

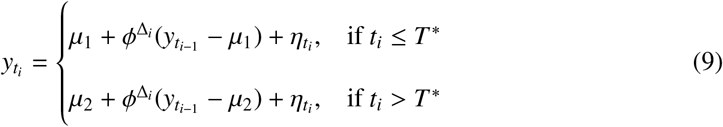

with 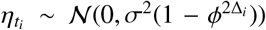. This formulation allows for both temporal dependence and a discrete regime change in the process mean.

We compare the performance of each model in terms of the mean squared error (MSE). To compute it, for a given rat, let 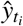 denote the point estimate of the duration of the nose press 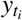 at a given time *t*_*i*_ and let *N* denote the number of sample points. Then the MSE is given by:

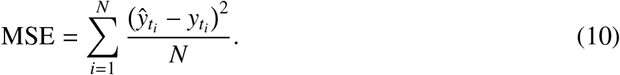

For each considered model, we can compute (10) and assess its performance, where smaller errors are preferred to order better models.

A small number of observations from Rats 7 and 10 were excluded from the model fitting due to their extreme values. For Rat 7, the final observation was excluded because it deviated substantially from the animal’s preceding measurements, being the largest value observed in the entire dataset for this animal, suggesting a non-representative artifact. For Rat 10, the rat that produced the largest number of responses in the session (1671 responses), seventeen observations –those that exceeded the 99th percentile– of that animal’s distribution were excluded to reduce the influence of extreme values. These exclusions accounted for less than 1.02% of the total observations.

## 3. Results

We fitted each of the models described in Section 2.1 to the time series of individual rats. The corresponding results are summarized in Table 1, where each row represents a rat and each column corresponds to one of the alternative model specifications introduced in the preceding section. Each cell reports the mean squared error (MSE) obtained by fitting the respective model to the series of a given rat, as defined in (10).

**Table 1:** MSE of fitted models per rat (ordered by increasing average MSE)

| Rat | CP | Log + iAR(1) | Log | Linear | Sqrt | Harmonics | iAR(1) + CP | iAR(1) | iAR(1)-γ | Exp |
| --- | --- | --- | --- | --- | --- | --- | --- | --- | --- | --- |
| 3 | **0.676** | 0.682 | 0.691 | 0.696 | 0.693 | 0.761 | 0.735 | 0.791 | 1.084 | 0.762 |
| 4 | **1.455** | 1.536 | 1.537 | 1.541 | 1.538 | 1.662 | 1.472 | 1.653 | 1.650 | 1.706 |
| 5 | **0.622** | 0.651 | 0.659 | 0.671 | 0.665 | 0.689 | 0.664 | 0.729 | 0.960 | 0.749 |
| 6 | **0.670** | 0.674 | 0.675 | 0.678 | 0.676 | 0.706 | 0.679 | 0.799 | 0.825 | 0.770 |
| 7 | **1.000** | 1.024 | 1.025 | 1.040 | 1.033 | 1.051 | 1.008 | 1.135 | 1.291 | 1.147 |
| 8 | 0.946 | 0.944 | 0.954 | **0.935** | 0.936 | 1.093 | 0.967 | 1.130 | 1.581 | 1.016 |
| 9 | **3.942** | 4.018 | 4.008 | 4.025 | 4.009 | 4.186 | 4.015 | 4.193 | 4.182 | 4.601 |
| 10 | **1.672** | 1.718 | 1.718 | 1.718 | 1.719 | 1.722 | 1.704 | 1.752 | 2.045 | 1.853 |

In nearly all cases, the structural changes model achieved the lowest level of mean squared errors (MSE). Simpler functional forms, such as linear, logarithmic, or square-root trends, provide adequate baseline fits, but consistently underestimate local variability: compared to a single changepoint in the mean, their errors are larger. The simple iAR with both gamma or Gaussian errors provides worse fits.

This is also noticeable when one compares the relative MSE in the heatmap of Figure 2 and in the scatter plot of Figure 3, where the change point specification systematically provides the minimum error, with all other models showing positive percent differences, with the exception being Rat 8. The relative excess error ranged from modest for most cases (1%-5%) to larger ones for rats with pronounced behavioral changes. Overall, these results indicate that the segmentation of data by a simple changepoint captures critical variance components that purely autoregressive structures or other functional forms fail to model.

**Figure 2:**
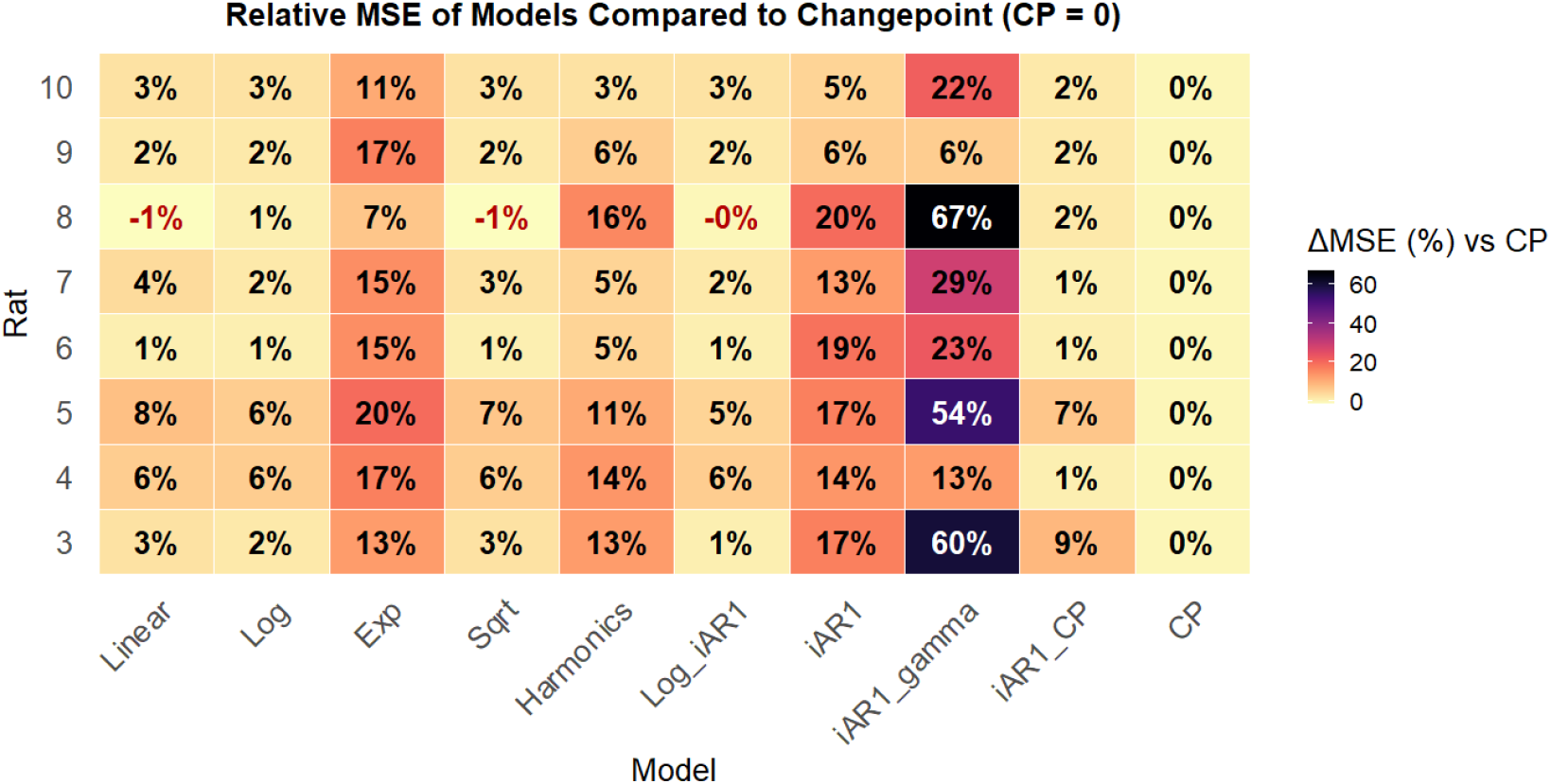
Relative mean square error (MSE). All models were compared to the change point (CP) model, considered as a reference, reason why the difference for all rats is null (0%).

**Figure 3:**
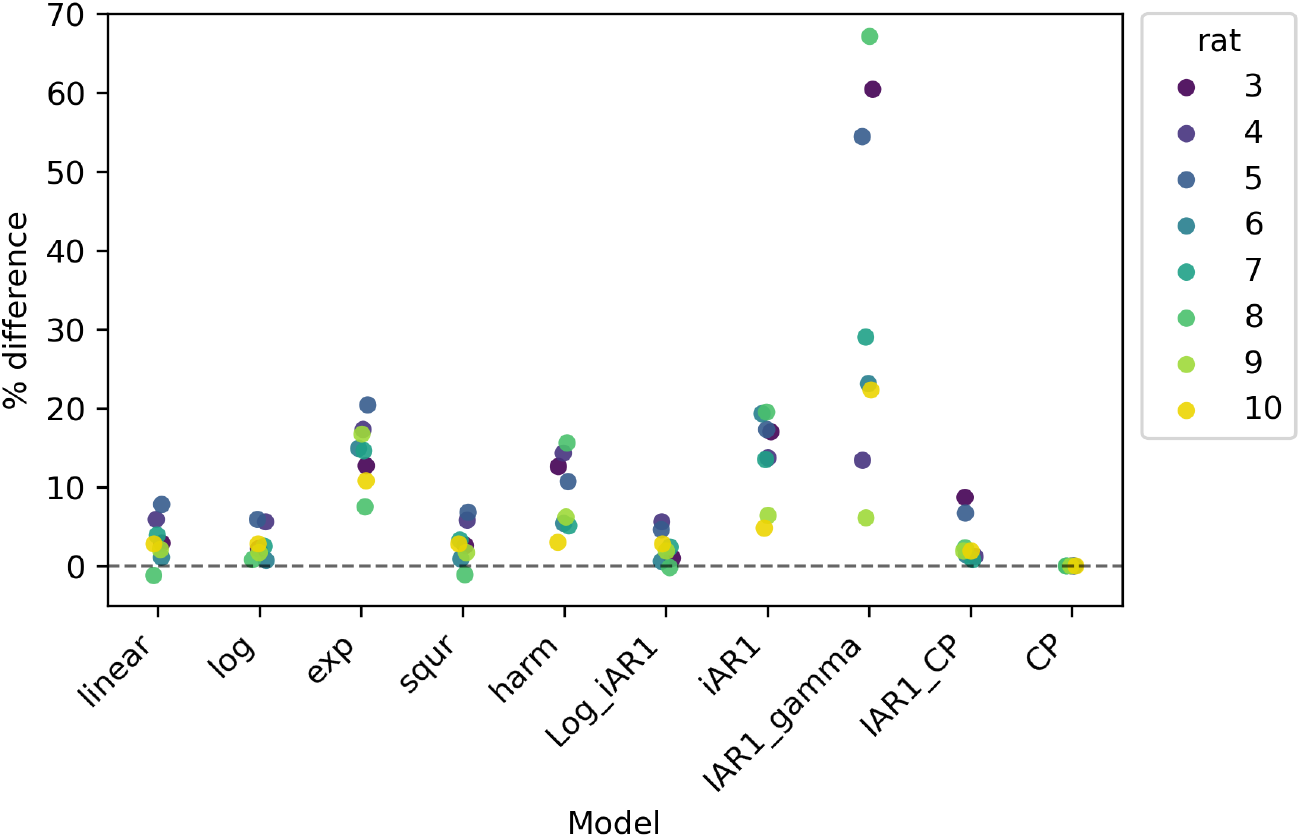
Percent difference of the MSE normalized by the values obtained with the changepoint model for each rat.

Given the previous results, the data suggests that a simple change in the mean better explains the dynamics of the response durations of the experiments. Due to that, we report in Table 2 the change points identified for each rat. We illustrate the different means computed prior to and after the break point, as well as the identified break in series, in Figure 4. Each panel depicts the observed duration series for an individual rat as in Figure 1, together with the fitted mean process obtained from the changepoint model as a solid red line.

**Table 2:** Change points identified for each rat.

| Rat | Break position | Breakpoint Time | Start time | End time | CPT time - start time | PCT OC |
| --- | --- | --- | --- | --- | --- | --- |
| 3 | 134 | 4485.15 | 3837.22 | 8395.11 | 647.93 | 14.22% |
| 4 | 212 | 4790.64 | 3735.22 | 7185.28 | 1055.42 | 30.59% |
| 5 | 268 | 5056.26 | 3819.75 | 11231.61 | 1236.51 | 16.68% |
| 6 | 223 | 4910.46 | 3995.86 | 8583.59 | 914.59 | 19.94% |
| 7 | 157 | 1245.82 | 242.27 | 14726.99 | 1003.54 | 6.93% |
| 8 | 482 | 3363.85 | 146.12 | 10433.83 | 3217.73 | 31.28% |
| 9 | 737 | 6997.16 | 65.33 | 14162.46 | 6931.84 | 49.17% |
| 10 | 116 | 707.52 | 105.78 | 13673.66 | 601.74 | 4.44% |

**Figure 4:**
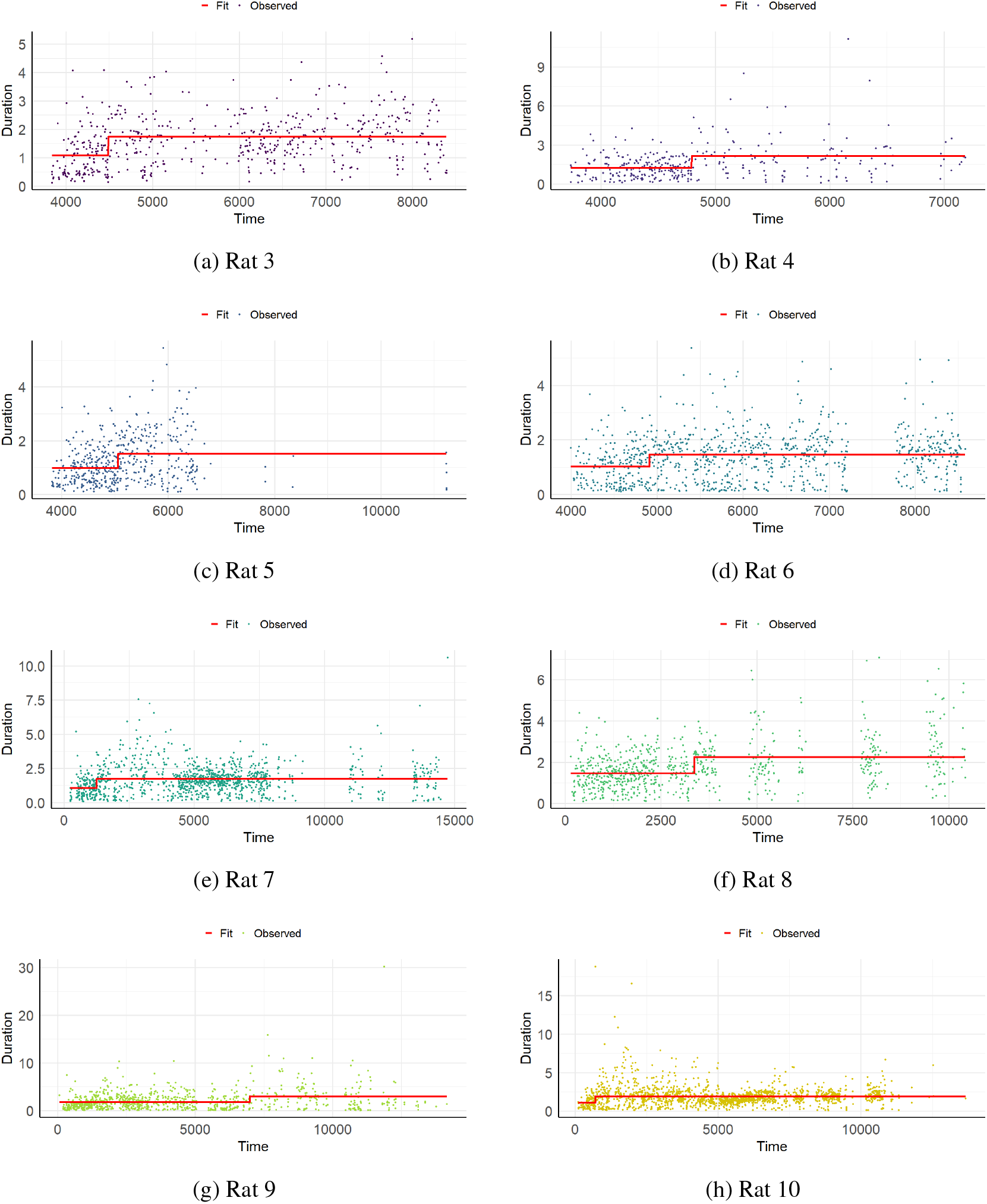
Modelling changepoints for durations

Across all subjects, the changepoint specification successfully identifies abrupt shifts in the level of the process, capturing episodes of structural modifications that are visually evident in the raw data. In all cases, the fitted mean increases after the breakpoint, which is coherent to the experiment: after some repetition, it is expected that the rats will understand that short nose pokes in the sensor will not produce rewards.

As reported in Table 2, the timing of the breakpoints is predominantly concentrated in the first half of the experiment, with 62.5% (5 out of 8 rats) occurring before 30% of the elapsed time and an additional 25% (2 out of 8 rats) between 30% and 32% of time elapsed. The only exception is Rat 9, whose changepoint occurs approximately midway through the experiment.

To evaluate whether the residuals of the change point specification account for the entire dependence structure in the data, we calculated Lomb-Scargle Periodogram (LSP) as of [16, 22] for each rat in Figure 5. As mentioned in Section 2.1, this spectral method is particularly suitable for unevenly spaced time series, as it generalizes the Fourier transform to irregularly sampled data. Overall, the LSP does not show peaks, suggesting that the residuals have no periodicity.

**Figure 5:**
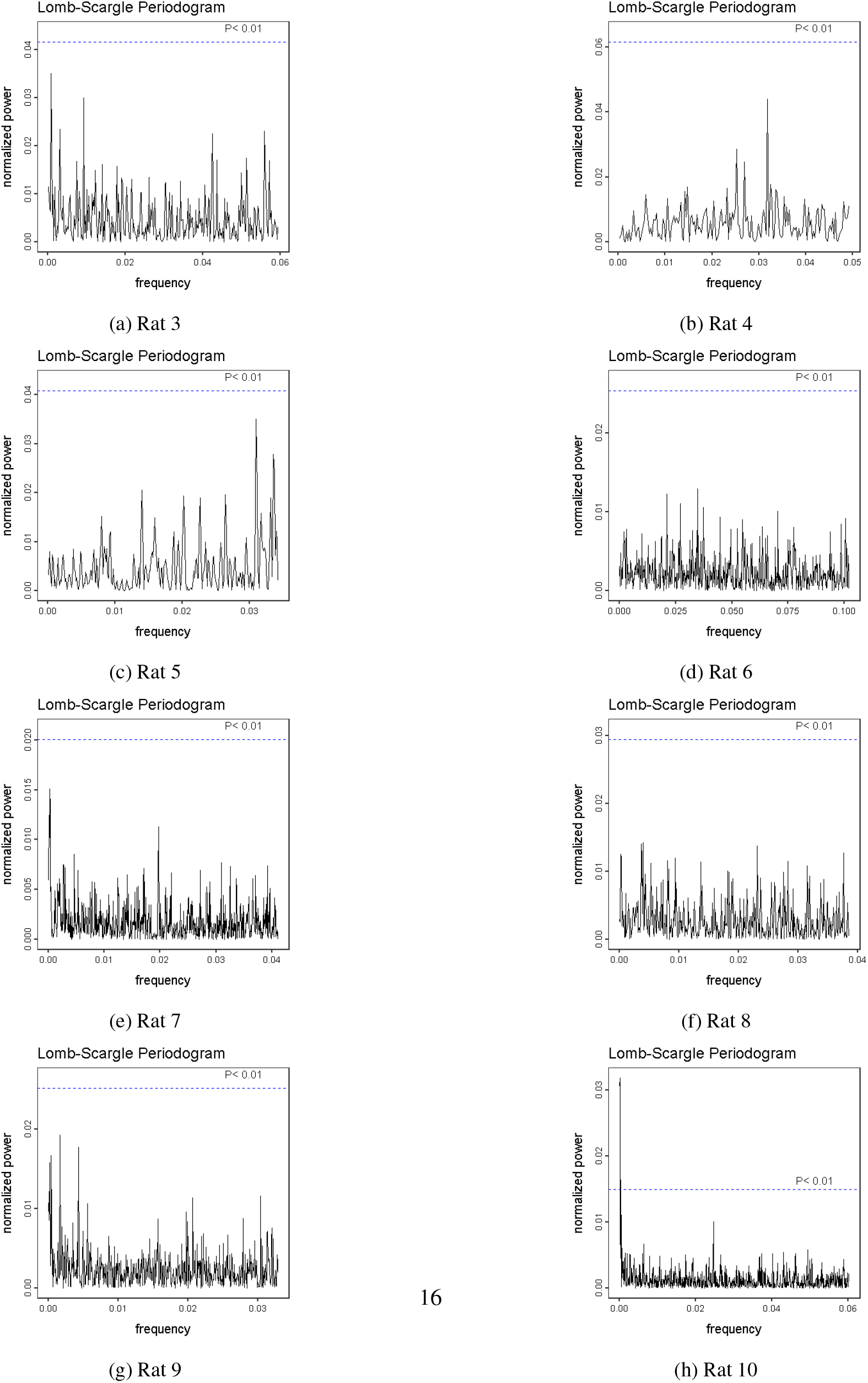
Lomb-Scargle Periodogram of the residuals

Overall, the results suggest that a simple changepoint model is better suited to explain the dynamics of the series. The residuals of the model do not seem to be statistically significantly correlated, suggesting a white noise-like behavior.

## 4. Discussion

The central objective of this study was to decide between two competing hypotheses regarding learning dynamics: a gradual, continuous process versus an abrupt, “snap-like” transition. Our results, summarized in Table 1 and Figure 3, provide robust support for the latter hypothesis within the context of the DRRD task. The consistent superiority of the single change-point (CP) model in seven of the eight animals, as measured by the lowest Mean Squared Error (MSE), strongly suggests that temporal learning in this paradigm is not best described by a smooth, continuous learning curve. Instead, the data are more consistent with a phase transition or a regime shift in behavior. The rats appear not to be gradually refining their response durations, but rather discovering and implementing a new behavioral strategy (holding the nose poke for longer) at a particular moment.

Our results align with the literature proposing “single-trial” or “Eureka-effect” learning [11, 20, 2], which is often obscured by analytical methods that aggregate data across trials or subjects. Figure 4 visually illustrates this transition, showing a consistent increase in the mean response duration after the identified breakpoint. This is biologically coherent: the animals learn that short-duration responses (below the 1.5s criterion) are unreinforced and subsequently shift their strategy to a new, longer-duration “set point.” Notably, Table 2 indicates that for the vast majority of subjects (7 out of 8), this change-point occurred relatively early in the experiment (five rats before 30 per cent of elapsed time and two just after 30 per cent). This suggests that the fundamental learning of the task rule can happen quickly, even if performance variability (the spikes and troughs in Figure 1) remains high after this shift.The sole exception was Rat 8, for which the simple linear model yielded a marginally lower MSE than the change-point model (0.935 vs. 0.946). This may indicate individual heterogeneity in learning strategies; perhaps for this specific animal, the process was indeed more gradual, or the high variability in its data (as seen in Figure 4f) obscured a clear breakpoint, allowing the linear model to provide a slightly better, albeit simplistic, fit.The analysis of the change-point model’s residuals, presented in Figure 5, is particularly telling. The use of the Lomb-Scargle Periodogram (LSP)—a spectral method appropriate for our unevenly spaced data—showed that after removing the abrupt shift in the mean, the residuals behave essentially as white noise (i.e., a flat spectrum with no significant peaks). This confirms that the simple change-point model was remarkably effective at capturing the entire temporal-dependence and learning structure in the data, rendering more complex autoregressive (like iAR(1)) or functional trend models unnecessary.This study also validates the proposed methodology. By treating operant behavior as a continuous, irregular time series rather than a sequence of discrete trials, we were able to apply statistical tools that honor the true temporal structure of the data. The traditional approach of “averaging by trial blocks” would have smeared this abrupt jump, artifactually making the learning appear gradual.While our results are robust, some limitations must be considered. We only tested for the presence of a single change-point. It is plausible that learning occurs in multiple, distinct stages, or that animals “forget” and “re-learn” the rule, which would be better modeled by multiple change-points. Furthermore, while our continuous-time approach implicitly incorporated Inter-Trial Intervals (ITIs) into the temporal structure (as Δ_*i*_), we did not actively model the ITI dynamics themselves, which, as noted in the introduction, may contain valuable information about motivation and strategy.

## 5. Final remarks

In this work, we investigated the temporal dynamics of learning in a Differential Reinforcement of Response Duration (DRRD) task. By abandoning traditional trial-based analysis and instead modeling behavior as a continuous and irregularly spaced time series, we were able to directly compare models of gradual versus abrupt learning.

Our results demonstrate robustly that a model incorporating a single, abrupt shift in the mean level of response durations (a “change-point”) explains the data variability significantly better than models assuming continuous change (linear, logarithmic) or simple autoregressive structures. The spectral analysis of this model’s residuals, via the Lomb-Scargle Periodogram, confirmed that the change-point model successfully captured the core learning structure, leaving behind uncorrelated white noise.

Collectively, these findings lend strong support to the hypothesis that, in this timing task, the fundamental learning process resembles an abrupt “insight” or phase shift rather than a slow, gradual refinement. This work validates the use of continuous-time series methods for analyzing operant behavior, providing a path to uncover the underlying dynamics of learning that are often masked by traditional averaging and trial segmentation techniques.

## Declarations

### Ethics approval

All procedures were approved by the Institutional Animal Care and Use Committee of the Federal University of ABC (CEUA-UFABC; permits 001/2014 and 020/2015).

### Consent to participate

Not applicable.

### Consent for publication

Not applicable.

### Data availability

The data underlying the results reported in this article are publicly available through the Open Science Framework at https://osf.io/uh4s6/.

### Code availability

The code used for the analyzes reported in this article is available from the corresponding author upon request.

